# The anti-tubercular activity of simvastatin is mediated by cholesterol-dependent regulation of autophagy via the AMPK-mTORC1-TFEB axis

**DOI:** 10.1101/2020.03.04.977579

**Authors:** Natalie Bruiners, Noton K. Dutta, Valentina Guerrini, Hugh Salamon, Ken D. Yamaguchi, Petros C. Karakousis, Maria L. Gennaro

**Author notes:** Corresponding author: Public Health Research Institute, New Jersey Medical School, Rutgers University, 225 Warren Street, Newark, NJ 07103.

## Abstract

Statins, which inhibit both cholesterol biosynthesis and protein prenylation branches of the mevalonate pathway, increase anti-tubercular antibiotic efficacy in animal models. We investigated the mechanism of anti-tubercular action of simvastatin in *Mycobacterium tuberculosis*-infected human monocytic cells. We found that the anti-tubercular activity of statins was phenocopied by cholesterol-branch but not prenylation-branch inhibitors. Moreover, statin treatment blocked activation of mechanistic target of rapamycin complex 1 (mTORC1), activated AMP-activated protein kinase (AMPK) through increased intracellular AMP:ATP ratios, and favored nuclear translocation of transcription factor EB (TFEB). These mechanisms all induce autophagy, which is anti-mycobacterial. The biological effects of simvastatin on the AMPK-mTORC1-TFEB-autophagy axis were reversed by adding exogenous cholesterol to the cells. Overall, our data demonstrate that the anti-tubercular activity of simvastatin requires inhibiting cholesterol biosynthesis, reveal novel links between cholesterol homeostasis, AMPK-mTORC1-TFEB axis, and intracellular infection control, and uncover new anti-tubercular therapy targets.

## 1. INTRODUCTION

As our knowledge of host-pathogen interactions grows for many infectious agents, pharmacologically manipulating the host response has emerged as a key approach to control or treat disease-causing infections. This approach may be best suited for infections caused by intracellular pathogens, given the intimate relationship between these pathogens and their host cells. For example, *Mycobacterium tuberculosis*, the intracellular pathogen causing tuberculosis, can subvert and disable the antimicrobial mechanisms of the host macrophage while adapting to the environmental conditions created by these mechanisms (1, 2). The treatment of active tuberculosis, which typically presents with lung tissue damage, is prolonged and requires multiple drugs, presumably due to the presence of phenotypically diverse *M. tuberculosis* subpopulations that exhibit antibiotic tolerance and poor penetration of antibiotics into infected tissues (3-5). The prolonged duration of anti-tubercular antibiotic therapy (a minimum of six months for drug-susceptible, uncomplicated tuberculosis) poses logistical difficulties for treatment providers and leads to poor patient compliance, particularly in low-resource regions (6). A dramatic consequence of these challenges is the development of antibiotic resistance, which requires treatments that are even more prolonged, less effective, more expensive, and more toxic (7).

One strategy that can address the above challenges is utilizing adjunctive chemotherapies that modify host responses to infection to reduce inflammation and tissue damage and/or to promote infection clearance (8). These host-directed therapies may shorten treatment duration, combat antibiotic-resistant disease by helping reduce its incidence, and improve treatment success, since it is unlikely that cross-resistance develops between antibiotics and host-directed therapeutics. The renewed attention to host-directed therapeutics warrants elucidating their mechanism of action in the context of the target infectious disease. Among the current drugs under evaluation as host-directed therapeutics against tuberculosis are statins (9-13). These drugs, which are prescribed worldwide as cholesterol-lowering agents, act by competitively inhibiting 3-hydroxy-3-methylglutaryl-CoA (HMG-CoA) reductase, the rate-limiting enzyme of the mevalonate pathway (14). This pathway regulates biosynthesis of cholesterol and isoprenoids, such as farnesyl pyrophosphate and geranylgeranyl pyrophosphate. The latter molecules are needed for protein prenylation, a process that activates and targets to membranes several protein classes that regulate cell growth, differentiation, and cell function (15). Consequently, statins not only possess cholesterol-lowering activity, but also exhibit pleotropic effects, such as tissue remodeling, inhibition of vascular inflammation, cytokine production, and immunomodulation (16, 17).

Due to their pleotropic effects, statins are currently under investigation in many pathological contexts. For example, their use has been explored against multisystem microbial infections, such as sepsis and pneumonia, and in non-infectious pathologies, such as autoimmune and inflammatory diseases (18-20). In regard to tuberculosis, statins exhibit anti-tubercular activity in murine and human macrophages infected *ex vivo* (9, 12, 13), display immunomodulatory effects in *ex vivo* infected human peripheral blood cells (13), and shorten the duration of antibiotic treatment in *M. tuberculosis*-infected mice (10, 11). Previous work (9) found a reduction in *M. tuberculosis* burden in murine macrophages by high doses of statins that inhibited both the cholesterol- and isoprenoid-forming branches of the mevalonate pathway. These effects were ascribed to the induction of autophagy, which controls *M. tuberculosis* infection (21-23), through the inhibition of geranylgeranyl biosynthesis, as observed in coronary arterial myocytes and certain cancer cell lines (24, 25). The molecular mechanisms underlying these results and the significance of effects induced by statin doses that far exceed therapeutic dosing remained unexplained. By utilizing simvastatin to treat *M. tuberculosis*-infected human monocytic cells, we now show that the anti-tubercular activity of statins results from their cholesterol-lowering effects and the consequent regulation of the nutrient- and energy-sensing pathways regulated by the AMP-activated protein kinase (AMPK), the mechanistic target of rapamycin complex 1 (mTORC1), and the transcription factor EB (the AMPK-mTORC1-TFEB axis), in ways that promote autophagy and control of infection with intracellular pathogens.

## 2. RESULTS

### 2.1. Simvastatin limits *M. tuberculosis* infection by inhibiting *de novo* synthesis of cholesterol

The mevalonate pathway, which is the mechanistic target of statins, is multi-branched (Fig. 1A). Identifying the pathway branch(es) associated with the observed anti-tubercular activity of these compounds is the first step toward elucidating the underlying mechanism of action. Treatment of *M. tuberculosis*-infected THP1 cells (a human monocytic cell line) with simvastatin, demonstrated that this drug decreased *M. tuberculosis* intracellular burden (Fig. 1 B), consistent with previous reports (11, 12). The anti-mycobacterial effect of simvastatin varied with the dose, with concentrations of 50-100 nM being the most effective (Fig. 1B). Additional experiments determined that these anti-mycobacterial doses were non-toxic for THP1 cells (supplementary Fig. 1A) and that simvastatin had no direct antimicrobial activity against *M. tuberculosis* in axenic cultures (supplementary Fig. 1B). We next tested the effect on intracellular *M. tuberculosis* burden of inhibitors specifically targeting each branch of the mevalonate pathway. As shown in Fig. 1A, the mevalonate pathway branches at farnesyl pyrophosphate (FPP) synthesis, which is used for protein prenylation, including farnesylation and geranylgeranylation, or is converted to squalene, which is required for *de novo* cholesterol synthesis. We treated *M. tuberculosis*-infected THP1 cells with inhibitors of farnesyl transferase (FTI-277), geranylgeranyl transferase - type 1 (GGTI-298) and 7-dehydrocholesterol reductase (DHCR7) (BM 15766 sulfate) (colored boxes in Fig. 1A). Moreover, since no inhibitor of geranylgeranyl transferase type 2 (GGTase-II) is commercially available, we used siRNA against the gene encoding the Rab escort protein 1 (REP-1), a chaperone protein that presents Rab proteins for prenylation by GGTase-II. We found that only cells treated with the cholesterol-branch inhibitor BM 15766, but not other branch-specific inhibitors, showed a reduced mycobacterial burden (35% inhibition relative to solvent-treated cells) (Fig. 1C). In addition, the inhibition of intracellular *M. tuberculosis* growth by simvastatin was reversed by exogenous addition of water-soluble cholesterol (Fig. 1D). We verified the link between simvastatin anti-mycobacterial activity and blockage of the cholesterol biosynthetic branch of the mevalonate pathway, as treatment with *M. tuberculosis*-inhibiting doses of simvastatin (50-100 nM) (Fig. 1 B) reduced the amount of free cholesterol in THP1 cells (Fig. 1E) while inducing no detectable alteration of the cellular ability to prenylate proteins (sentinel targets of each prenylation pathway are shown in Fig. 1F). Taken together, these results clearly demonstrate that the anti-tubercular activity of simvastatin specifically targets the cholesterol biosynthetic branch rather than the prenylation branches of the mevalonate pathway.

**Figure 1.**
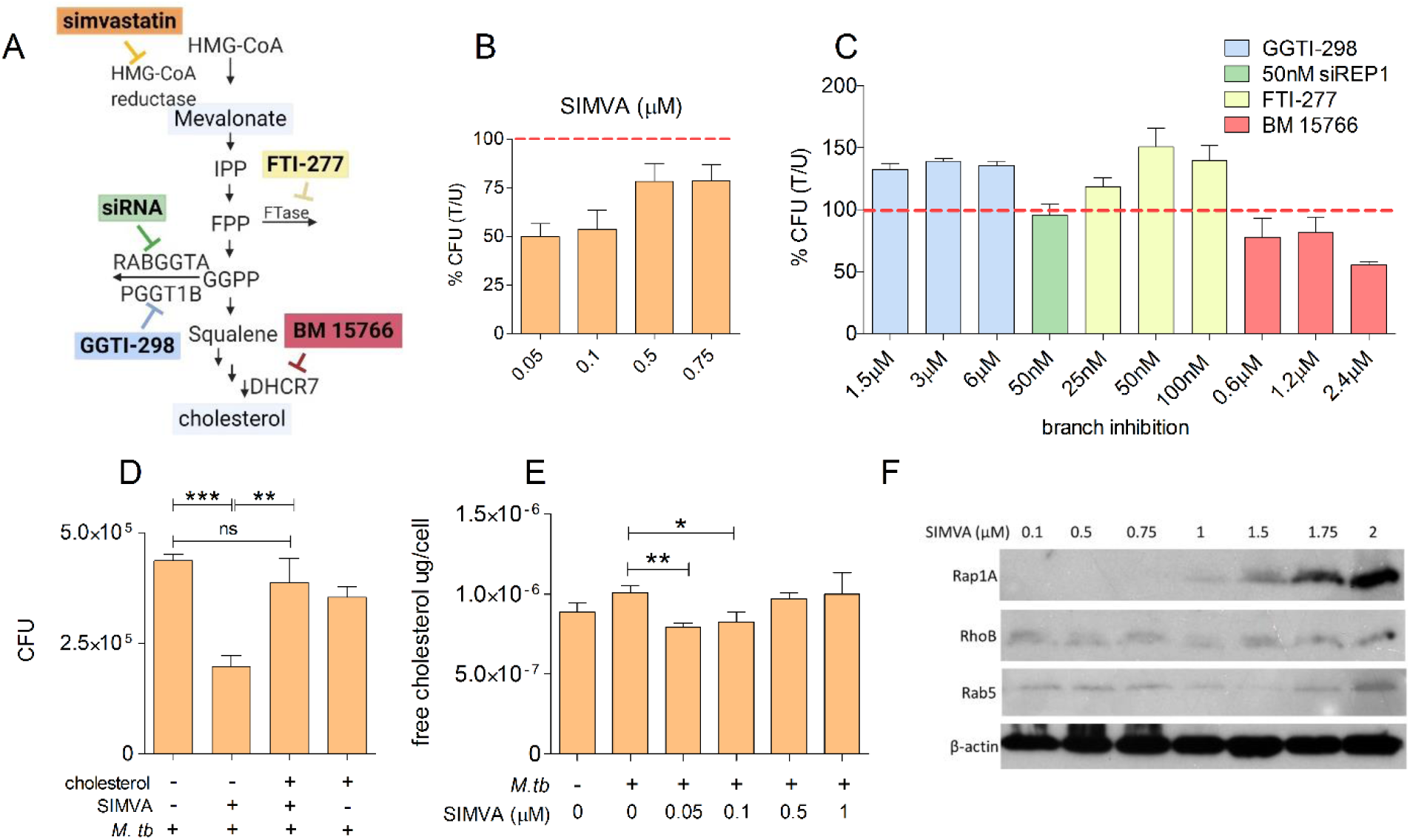
Simvastatin reduces the intracellular burden of *M. tuberculosis* by inhibiting cellular cholesterol biosynthesis. (**A**) Diagram of the mevalonate pathway. Abbreviations: HMG-CoA, 3-hydroxy-3-methylglutaryl-coenzyme A; IPP, isopentenyl pyrophosphate; FPP, farnesyl pyrophosphate; FTase, Farnesyltransferase; GGPP, geranylgeranyl pyrophosphate; RABGGTA, Rab Geranylgeranyltransferase Subunit Alpha; PGGT1B, Protein Geranylgeranyltransferase Type I Subunit Beta; DHCR7, 7-dehydrocholesterol reductase. The colored boxes represent pharmacological inhibitors of specific enzymes in the pathway (simvastatin, FTI-277, GGTI-298, BM 15766), as indicated. siRNA targeting REP-1 [a chaperone protein that presents Rab proteins for prenylation by geranylgeranyltransferase type II (GGTase-II)] was used, since no pharmacological inhibitor of GGTI type 2 was commercially available. (**B, C**) THP1 cells were infected with *M. tuberculosis* H37Rv, followed by treatment with multiple doses of simvastatin (**B**) or branch inhibitors of the mevalonate pathway (**C**). CFU was enumerated at 6 days post treatment. Data are presented as percent CFU changes relative to solvent control. U, solvent-treated; T, treated. The red dotted line represents *M. tuberculosis* CFU levels in the absence of any pharmacological treatment. (**D**) Analysis of cellular cholesterol levels in infected THP1 cells treated with solvent and increasing doses of simvastatin. (**E**) Intracellular growth of *M. tuberculosis* in THP1 cells treated for 6 days with 100 nM simvastatin in the absence and presence of water-soluble cholesterol (1.25 μg/mL). Significance was tested by Student’s t-test with n = 3 (*, p < 0.05, **, p < 0.01 and ***, p < 0.001). (**F**) Western blot analysis of sentinel protein targets of the mevalonate pathway of infected THP1 cells treated with simvastatin (SIMVA; 0.1 – 2 μM) for 6 days. Beta-actin was used as loading control.

### 2.2. Simvastatin inhibits *M. tuberculosis* growth by inducing autophagy in a cholesterol-dependent manner

Seminal findings by Segal and Bloch, which demonstrated that *M. tuberculosis* preferentially utilizes lipids as a carbon and energy source during infection (26), have spurred vast research efforts into host lipid requirements for survival of this pathogen during infection (27-29). In particular, it is known that *M. tuberculosis* metabolizes cholesterol during infection and that degradation of this sterol contributes to the pathogen’s survival in host cells (30, 31). Therefore we asked whether the reduction of intracellular *M. tuberculosis* burden by simvastatin was due to the reduced availability of host cholesterol as a nutrient source during host cell infection. Catabolism of cholesterol by beta oxidation produces propionyl-CoA, which is assimilated into pyruvate via the methyl citrate cycle (32). *M. tuberculosis* mutants that are defective in methyl citrate cycle enzymes cannot grow either in synthetic media containing cholesterol as a primary carbon source or in macrophages (29). Moreover, the intracellular growth defect of such mutants is relieved by additional mutations impairing mycobacterial cholesterol uptake (32), implying that accumulation of propionyl-CoA is toxic. Therefore, we predicted that pharmacologically induced reduction of host cell cholesterol should relieve the intracellular growth defect of methyl citrate cycle*-*deficient *M. tuberculosis* mutants.

To test our hypothesis, we used *M. tuberculosis* wild-type (WT) and a mutant strain genetically inactivated in *rv1129c*, which encodes a transcriptional factor required to induce the methyl citrate cycle genes (32, 33). When we infected THP1 cells with *M. tuberculosis* WT and Δ*rv1129c* mutant strains, we found that the mutant strain was attenuated for growth in macrophages, as expected (up to three-fold reduction of CFU relative to WT) (Fig. 2A, dark blue bars). Treating infected cells with BM 15766 sulfate (which specifically inhibits *de novo* cholesterol synthesis, Fig. 1A) drastically reduced intracellular growth of the WT strain (45%), as expected, but increased growth in the Δ*rv1129c* mutant strain by 50% (Fig. 2A). This result supports our initial hypothesis. In contrast, simvastatin treatment inhibited WT and mutant growth to a similar extent (>40% growth inhibition) (Fig. 2B). The different effect of the two drugs on mutant growth strongly implies that the reduced intracellular burden of *M. tuberculosis* in simvastatin-treated cells is not primarily (or solely) caused by the reduced availability of cholesterol as carbon source for mycobacterial growth.

**Figure 2.**
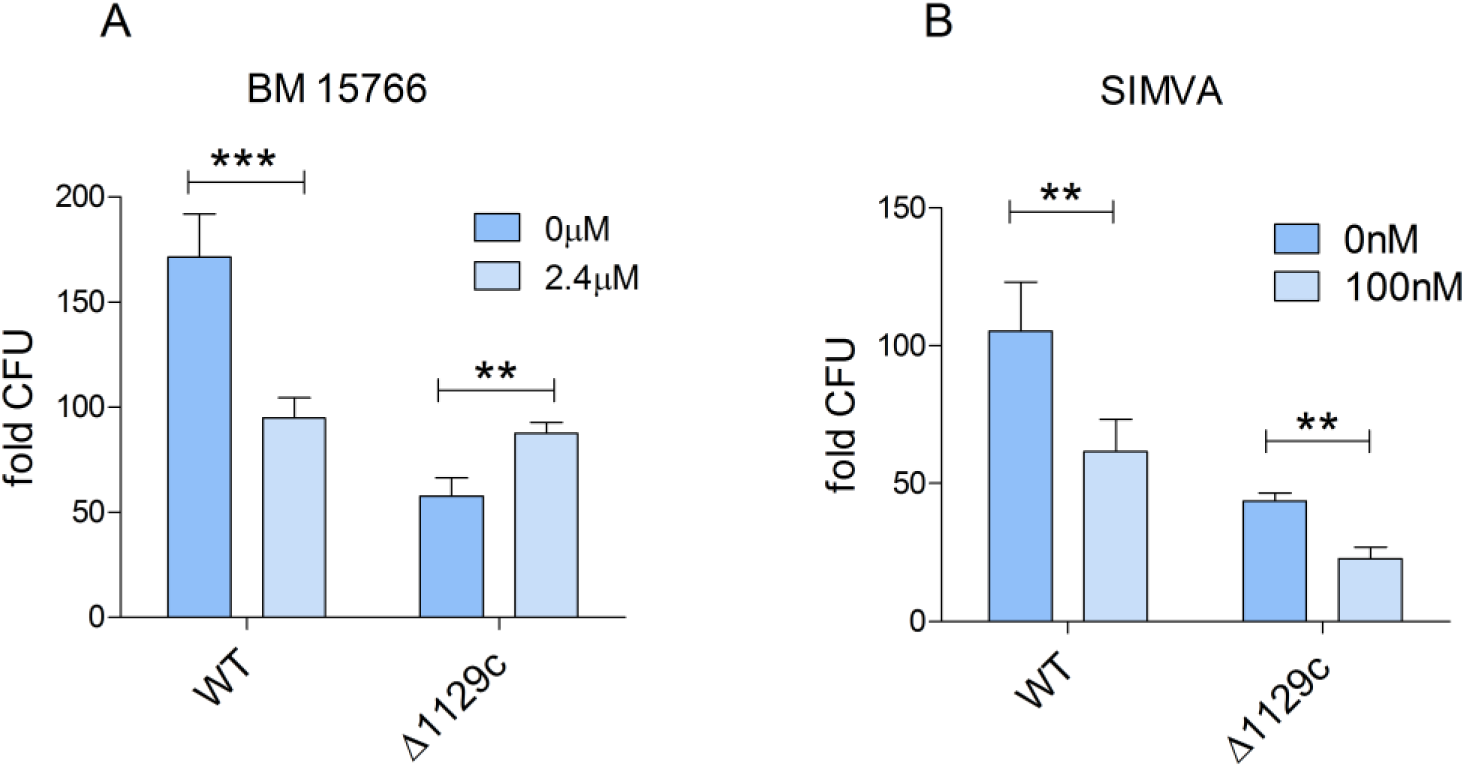
Reduction of *M. tuberculosis* growth by simvastatin is independent of mycobacterial utilization of cholesterol as a carbon source. (**A, B**) THP1 cells were infected with wildtype (WT) or deletion mutant (ΔRv1129c) *M. tuberculosis* strains for 6 days and treated with either BM 15766 (inhibitor of 7-dehydrocholesterol reductase) (2.4 μM) (**A**) or simvastatin (100 nM) (**B**). At day 6 post-treatment, cells were lysed, plated on Middlebrook 7H10 agar plates and incubated for 3 weeks at 37°C for bacterial CFU enumeration. Data represent fold change in CFU at 6 days relative to 4 hours post-infection. Here and in subsequent figures data are expressed as mean ± SD from biological triplicates. Significance was tested by Student’s t-test (**, p < 0.01 and ***, p < 0.001).

A previous report showed that simvastatin induces autophagy, a known anti-mycobacterial function (34), in *M. tuberculosis*-infected bone marrow–derived macrophages (9). That effect was observed with high drug doses (50 µM), well above those affecting all branches of the mevalonate pathway (Fig. 1F) and that are toxic to THP1 cells (supplementary Fig. 1A). We used simvastatin at 100nM concentration to examine whether this drug induces autophagy at a dose that is anti-mycobacterial (Fig. 1B) while only affecting cholesterol biosynthesis (but not protein prenylation) in THP1 cells (Figs. 1E and 1F). To assess the effects of simvastatin treatment on autophagy we measured the levels of the autophagy marker p62/sequestosome 1, which is reduced upon induction of autophagy and increased when autophagy is impaired (35). As expected, infection of THP1 cells with *M. tuberculosis* increased p62 levels (Figs. 3A and 3B), since *M. tuberculosis* infection inhibits autophagy (36). When we treated the infected cells with simvastatin, we found that p62 protein levels decreased (Figs. 3A and 3B), indicating activation of autophagy. The effect of simvastatin on p62 protein levels was reversed by the addition of exogenous cholesterol, demonstrating a role for cholesterol in autophagy regulation by statins (Figs. 3A and 3B).

**Figure 3.**
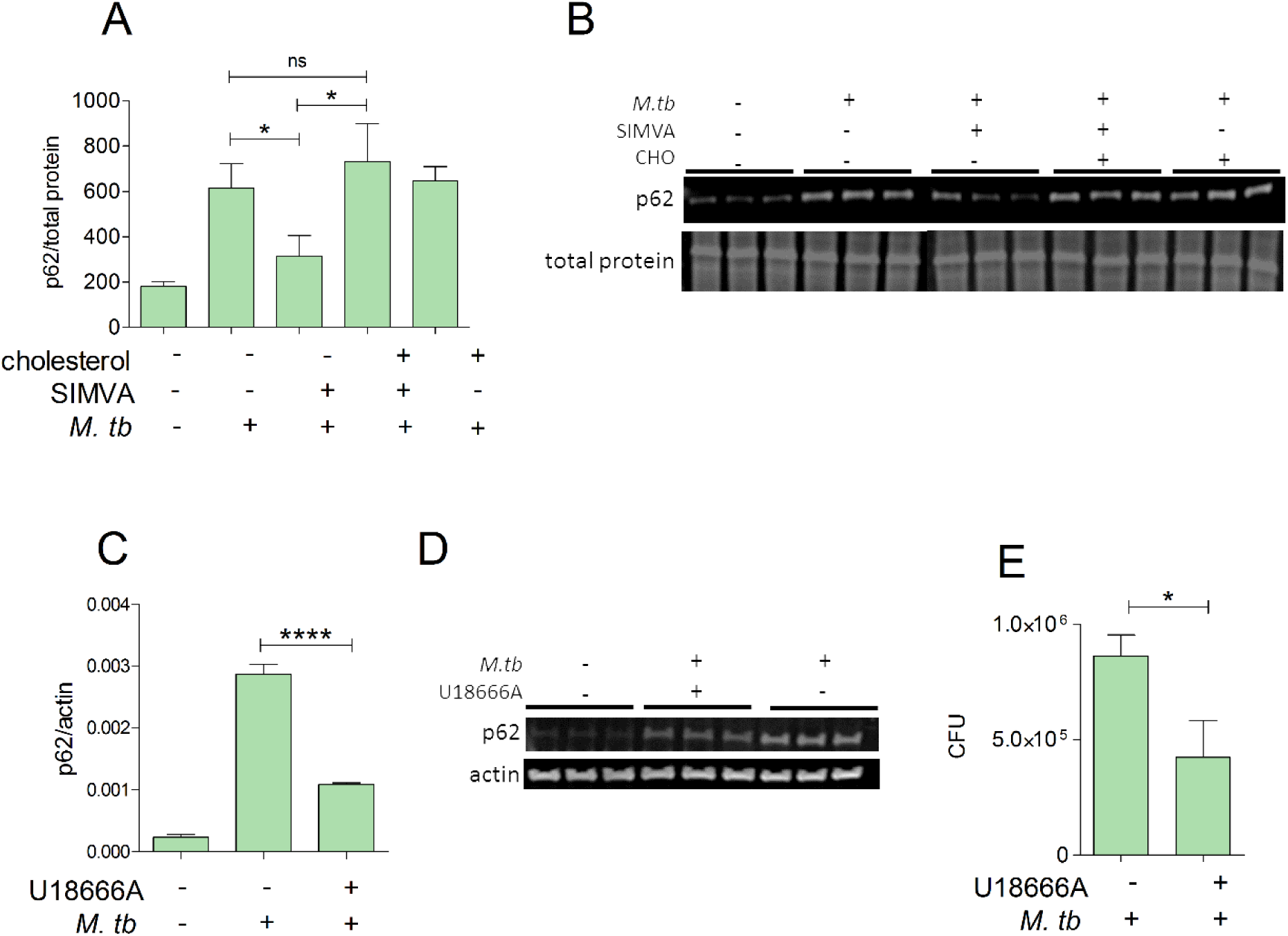
Reduction of *M. tuberculosis* growth by simvastatin is associated with cholesterol-dependent induction of autophagy. **(A, C)** Immunoblot analysis of the abundance of p62/SQSTM1 and beta-actin or total protein in whole-cell lysates obtained from THP1 cells infected with *M. tuberculosis* for 6 days and treated with DMSO as a solvent control or (**A**) 100 nM simvastatin in the absence or presence of water-soluble cholesterol (1.25 μg/mL) and (**C**) U186666A (inhibitor of cholesterol transport and synthesis) (1.25 μM). Protein quantification and normalization relative to total protein or beta-actin per lane, was performed using LI-COR Image Studio software. (**B, D**) Representative immunoblots of p62/SQSTM1 in whole cell lysates of THP1 cells treated with (**B**) 100nM simvastatin in the presence of soluble cholesterol (1.25μg/mL) and (**D**) U186666A (1.25 μM). When total protein was used as loading control (panel B), only a portion of the membrane probed for total protein is shown. (**E**) Effect of U18666A on intracellular growth of *M. tuberculosis* in THP1 cells. Significance was tested by Student’s t-test (*, p < 0.05 and ****, p < 0.0001); ns, not significant.

If interfering with cholesterol homeostasis modulates autophagy, treating infected macrophages with a class of drugs (other than statins) impacting cellar cholesterol homeostasis should also induce autophagy and have anti-mycobacterial activity. When we treated *M. tuberculosis*-infected THP1 cells with U18666A, which inhibits intracellular cholesterol transport (37), we observed a >60% reduction of p62 levels compared to infected, solvent-treated control cells (Figs. 3C and 3D). The autophagy effect was accompanied by anti-mycobacterial activity (>45% *M. tuberculosis* growth inhibition, Fig. 3E). Taken together, these results demonstrate that simvastatin induces autophagy by altering cellular cholesterol homeostasis.

### 2.3. Simvastatin induces autophagy by inhibiting mechanistic target of rapamycin complex 1 (mTORC1) activation in a cholesterol-dependent manner

A key inhibitor of autophagy is mTORC1, a serine-threonine protein kinase that functions as a nutrient/energy/redox sensor. Statins can alter mTORC1 signaling and autophagy in coronary arterial myocytes by affecting prenylation-dependent mechanisms (24). Therefore we tested whether simvastatin treatment at 100 nM, which only affects the cholesterol biosynthetic branch of the mevalonate pathway (Fig. 1C and 1F), inhibited mTORC1 activity in *M. tuberculosis*-infected cells. Since the enzymatic activity of mTORC1 is dependent on phosphorylation of the mTOR subunit, we measured phosphorylation at serine 2448, which is an mTORC1 activation mark (38-40), in infected THP1 cells treated with solvent control or 100 nM simvastatin. We found that *M. tuberculosis* infection increased mTORC1 activation, while simvastatin decreased it to uninfected levels (Figs. 4A and 4B). Further, simvastatin-mediated reduction of mTORC1 activation was reversed by supplementing the cell culture with water-soluble cholesterol (Figs. 4A and 4B). This result links the simvastatin effects on mTORC1 activity to cellular cholesterol homeostasis. To further establish the relationship between mTORC1 activity and simvastatin-mediated induction of autophagy, we supplemented simvastatin-treated cells with L-arginine, a known activator of mTORC1 (41). We found that adding L-arginine abolished the simvastatin-induced inhibition of *M. tuberculosis* burden (Fig. 4C), implying that mTORC1 affects mycobacterial burden. Indeed, treatment with everolimus, an allosteric inhibitor of the mTOR subunit of the mTORC1 complex (42, 43), reduced the *M. tuberculosis* burden of THP1 cells by 25% (Fig. 4D). Taken together, these results indicate that reduction of cellular cholesterol inhibits mTORC1 activity thereby inducing autophagy and, consequently, inhibiting *M. tuberculosis* infection.

**Figure 4.**
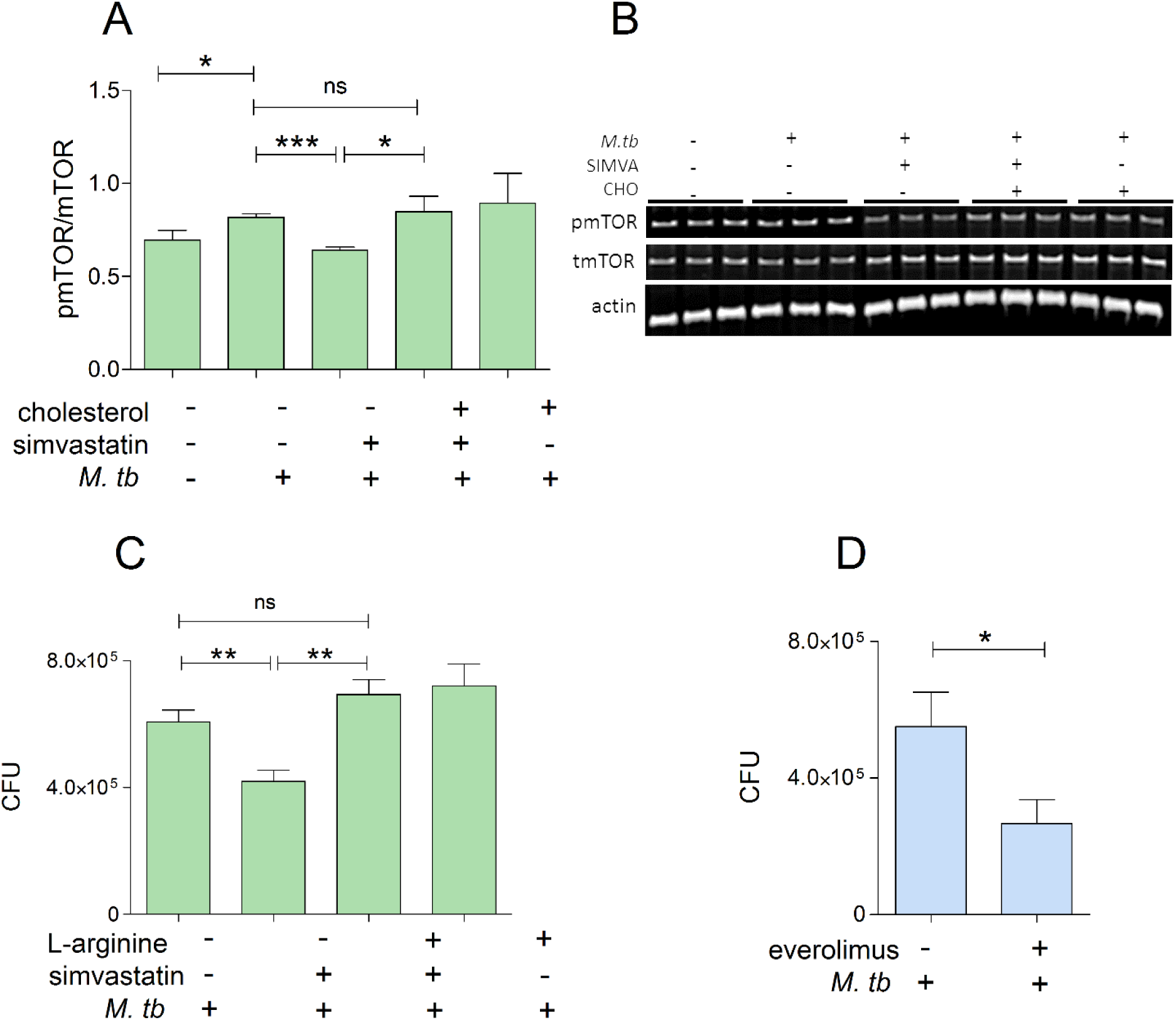
Reduction of *M. tuberculosis* growth by simvastatin is associated with cholesterol-dependent reduction of mTORC1 activation. (**A**) Immunoblot analysis of the phosphorylated/total mTORC1 ratio normalized to β-actin in whole-cell lysates of uninfected and infected THP1 cells for 6 days treated with DMSO as a solvent control or 100nM simvastatin in the absence and presence of soluble cholesterol (1.25 μg/mL). Protein quantification and normalization relative to beta-actin per lane, was performed using LI-COR Image Studio software. (**B**) A representative image of immunoblots as shown in **A**. (**C**) Intracellular growth of *M. tuberculosis* in THP1 cells infected for 6 days and treated with 100 nM simvastatin in the absence or presence of L-arginine (0.78 mM). (**D**) Effect of the mTORC1 inhibitor everolimus on intracellular growth of *M. tuberculosis* in THP1 cells. Significance was tested by Student’s t-test (*, p < 0.05, **, p < 0.01 and ***, p < 0.001)); ns, not significant.

### 2.4. Simvastatin regulates the nuclear translocation of transcription factor EB (TFEB)

We next sought to identify the mTORC1-dependent pathway(s) through which simvastatin induces autophagy and controls *M. tuberculosis* infection. A prime candidate is TFEB, a transcription factor involved in lysosomal biogenesis and autophagy induction (44, 45). mTORC1 inhibits the nuclear translocation of TFEB, which is required for its activation (46, 47). We found that simvastatin treatment increased TFEB abundance in the nuclear fraction but not in the total extracts obtained from *M. tuberculosis*-infected cells compared to infected control cells (exposed to solvent alone) (Fig. 5A). Exogenously added cholesterol reversed the simvastatin effect on TFEB nuclear localization (Fig. 5B). Moreover, treatment with digoxin, a TFEB activator, reduced the intracellular bacillary burden by 30% in infected THP1 cells (Fig. 5C). Taken together, these data imply that by controlling cholesterol levels, simvastatin induces nuclear localization and consequent activation of TFEB, an mTORC1-regulated transcription factor that induces autophagy.

**Figure 5.**
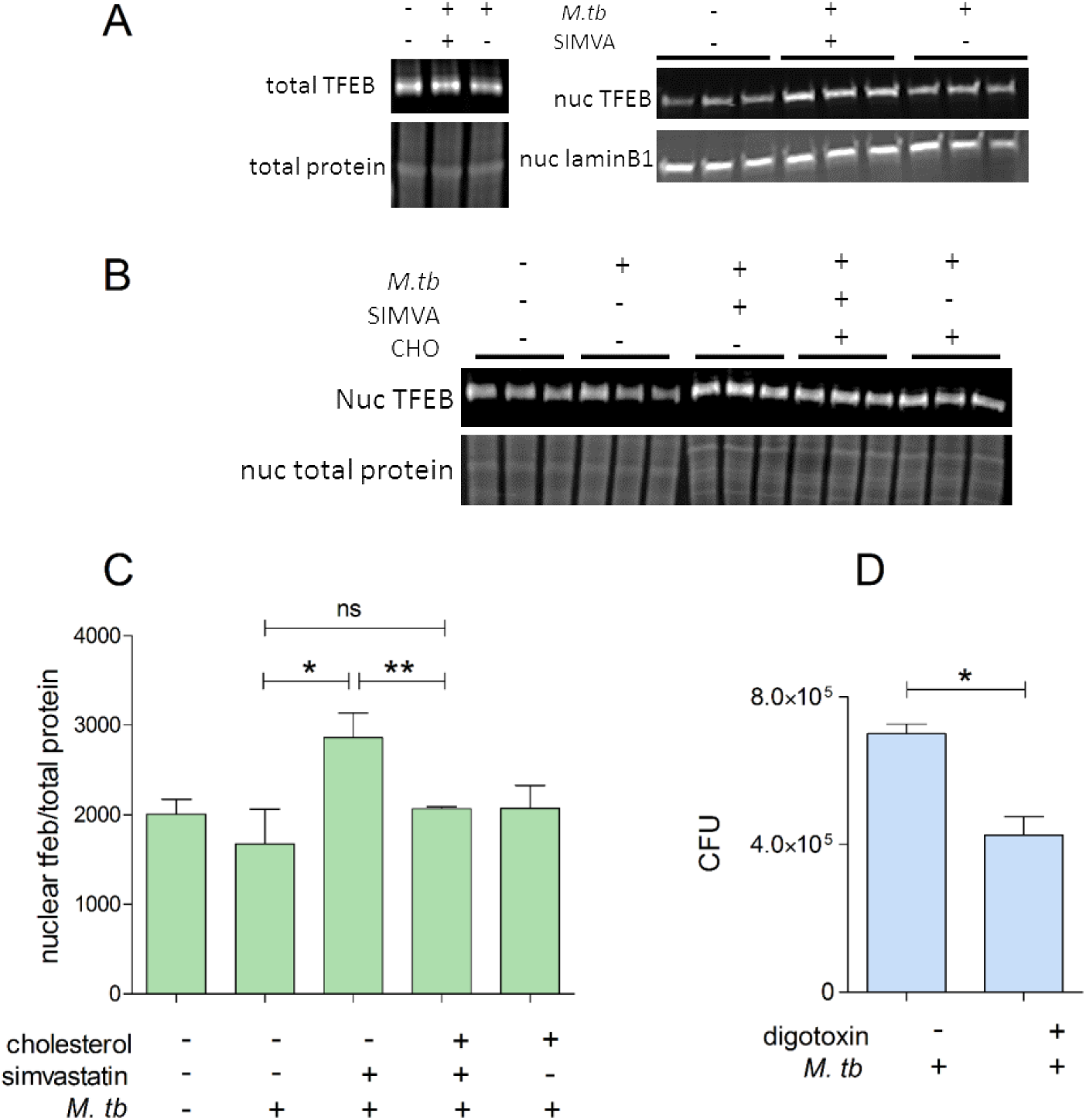
Reduced *M. tuberculosis* burden by simvastatin is associated with cholesterol-dependent nuclear translocation of TFEB. **(A)** Representative immunoblots of TFEB in whole-cell lysates (left panel) and nuclear extracts (right panel) of THP1 cells uninfected or infected with *M. tuberculosis* for 6 days treated with DMSO as a solvent control or 100nM simvastatin. Protein quantification and normalization relative to total protein or lamin B1, per lane, was performed using LI-COR Image Studio software. (**B**) Immunoblot analysis of TFEB abundance (relative to total protein) in nuclear lysates of infected THP1 cells treated with 100 nM simvastatin in the absence or presence of soluble cholesterol (1.25 μg/mL). When total protein was used as loading control (panel A and B), only a portion of the membrane probed for total protein is shown. (**C**) Effect of 7.5 nM Digoxin (TFEB activator) on the intracellular growth of *M. tuberculosis* in THP1 cells. Data are shown as means +/-SD from 3-4 wells. Significance was tested by Student’s t-test (*, p < 0.05); ns, not significant.

### 2.5. Simvastatin regulates transcription of the mTORC1 and TFEB pathways

We next sought corroboration of our THP1 mechanistic results in transcriptomics data generated with simvastatin-treated human primary cells. We searched for relevant molecular mechanisms by re-analyzing the results of a gene expression study of simvastatin-treated human peripheral blood mononuclear cells (PBMC) performed in the context of cardiovascular disease [(GSE4883) and (48)]. We found strong evidence that TFEB-regulated genes were upregulated by simvastatin treatment of PBMC (TFEB-bound in Table 1). Moreover, the mTOR signaling pathway was also upregulated (Table 1). The top three drivers of this result were genes encoding growth factor receptor-bound protein 10 (GRB10), unc-51 like autophagy-activating kinase 2 (ULK2), and sestrin 2 (SESN2) (supplementary Table 1). Upregulation of these three genes is associated with diminished mTORC1 activity, since GRB10 and SESN2 are negative regulators of the mTORC1 signaling pathway (49, 50), while ULK2, which promotes autophagy (51), is downregulated by mTORC1 (52). In addition, pathways expressing functions that require mTORC1 activation, such as ribosome biogenesis and hypoxia factor 1 alpha (HIF1A)-bound genes (HIF1A activity is induced by mTORC1 (53)), were reduced by simvastatin treatment, while insulin-receptor signaling, which is inhibited by mTORC1 (54), was found upregulated in simvastatin-treated cells. Thus, simvastatin treatment induced a clear transcriptomic signature of decreased mTORC1 activity and increased autophagy (Table 1).

**Table 1.** Transcriptomic changes of signaling pathways associated with mTORC1 and TFEB in simvastatin-treated PBMCs (GEO ID: GSE4883). Shown are pathways of interest ranked according to increased or decreased transcriptional changes that are statistically significant.

| Pathway | Rank | p value | FDR | Gene-set Source |
| --- | --- | --- | --- | --- |
| ↑ TFEB bound genes | 3 | 0.00038 | 0.00617 | ChIP |
| ↑ mTOR signaling | 13 | 5.82E-05 | 0.0047 | KEGG Pathways |
| ↑ IRS activation | 17 | 0.00011 | 0.00723 | Reactome Pathways |
| ↓ ribosome biogenesis | 21 | 8.34E-06 | 0.00034 | KEGG Pathways |
| ↓ HIF1A bound genes | 29 | 0.00053 | 0.0013 | ChIP |
| ↑ autophagy | 61 | 0.00159 | 0.01609 | KEGG Pathways |

### 2.6. Simvastatin induces activation of AMP-activated protein kinase (AMPK)

Our pathway analysis of simvastatin-treated PBMC also identified upregulation of signaling pathways regulated by AMPK, a serine-threonine kinase that inhibits mTORC1 signaling and induces autophagy (55) and upregulated Liver kinase B1 (LKB1), a serine-threonine kinase that activates AMPK (56) (Table 2), as previously observed in squamous cell carcinoma (57). Since these two proteins participate in the energy-sensing cascade activated by an increased AMP:ATP ratio (56), we first asked whether simvastatin treatment alters cellular AMP:ATP ratios in *M. tuberculosis*-infected cells. We did find that simvastatin reversed the infection-induced decrease in AMP:ATP ratio (Fig. 6A). In accord with these results, treating THP1 cells with compound C, an AMPK inhibitor (58), reversed the reduction of intracellular *M. tuberculosis* burden by simvastatin (Fig. 6B). Furthermore, treatment of infected THP1 cells with A-769662, an AMPK activator (59), reduced *M. tuberculosis* burden by 40% (Fig. 6C). Thus, simvastatin favors the anti-tubercular effect of AMPK by positively regulating both AMPK activation state through AMP:ATP levels and AMPK expression levels.

**Table 2.** Transcriptomic changes of signaling pathways associated with AMPK activation in simvastatin-treated PBMCs (GEO ID: GSE4883). Shown are pathways of interest ranked according to increased or decreased transcriptional changes that are statistically significant.

| Pathway | Rank | p value | FDR | Gene-set Source |
| --- | --- | --- | --- | --- |
| ↑ LKB1 signaling | 3 | 5.65E-06 | 0.00343 | Pathway Interaction Database |
| ↑ AMPK signaling | 69 | 0.0036 | 0.03063 | KEGG Pathways |

**Figure 6.**
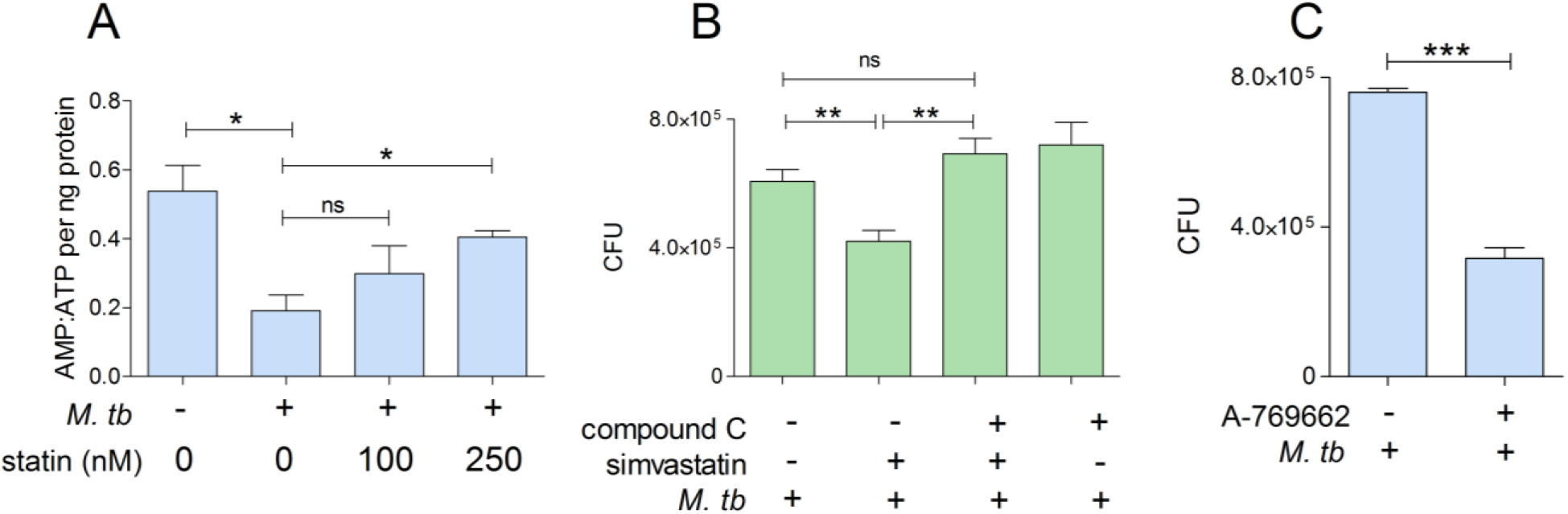
Reduction of *M. tuberculosis* growth by simvastatin is associated with regulation of the AMPK pathway. (**A**) AMP:ATP ratio in THP1 cells treated with 100 and 250 nM simvastatin. Abbreviations: statin, simvastatin. (**B**) Intracellular growth of *M. tuberculosis* in THP1 cells infected for 6 days and treated with DMSO as a solvent control or 100 nM simvastatin in the absence or presence of compound C (AMPK inhibitor) (110 nM). (**C**) Effect of 25 μM A-769662 (AMPK activator) on the intracellular growth of M. *tuberculosis* in THP1 cells.

## 3. DISCUSSION

Here we show that inhibition of the cholesterol biosynthesis branch of the mevalonate pathway underlies the anti-tubercular activity of statins. Simvastatin reduces *M. tuberculosis* burden in human macrophages by decreasing cholesterol levels, thereby regulating the AMPK-mTORC1-TFEB axis in ways that promote autophagy, which is anti-mycobacterial. The anti-tubercular effects of statin-mediated cholesterol reduction occur by inducing autophagy rather than by limiting access to cholesterol as a carbon source for intracellular mycobacteria. Moreover, statin treatment of human cells has transcriptional effects on mTORC1 and AMPK signaling pathways. Together, these findings reveal the hitherto unknown mechanistic foundations of the anti-tubercular activity of statins (diagrammed in Fig. 7). Our results support the ongoing testing of statins as adjunctive therapy against tuberculosis (60) and identify novel cellular targets for additional host-directed anti-tubercular treatments.

**Figure 7.**
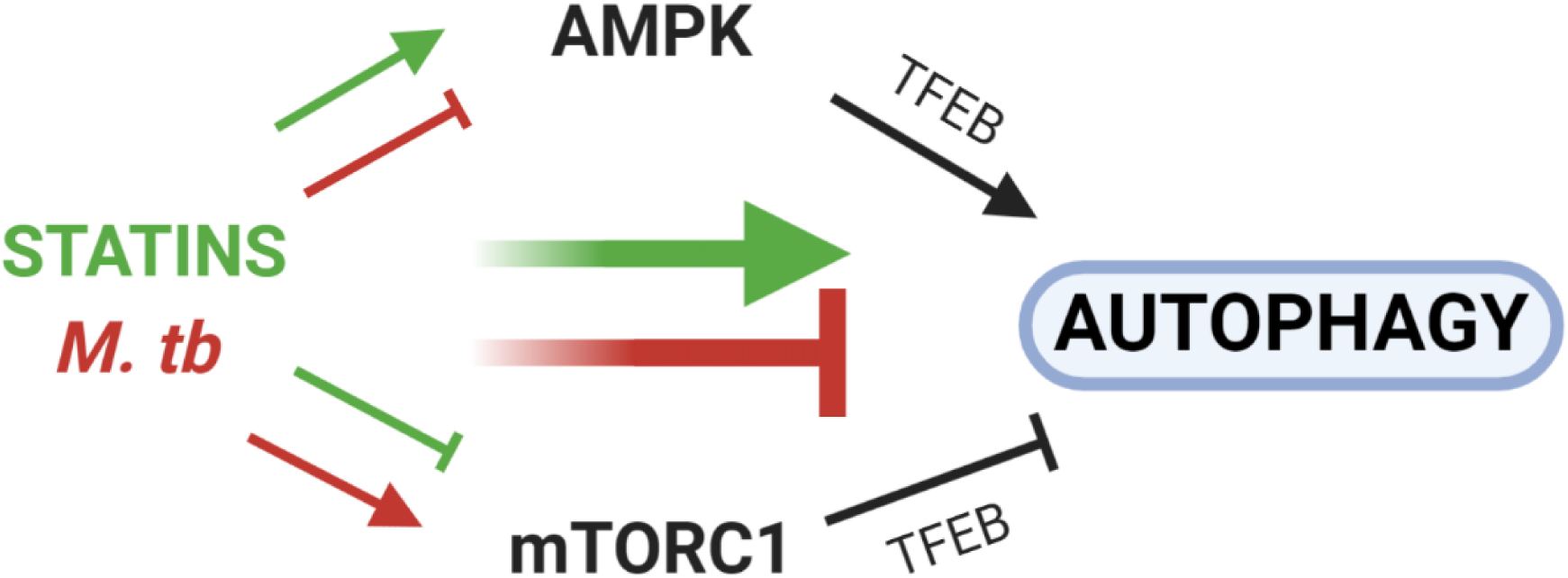
Statins and *M. tuberculosis* infection exhibit opposing effects on AMPK, mTORC1, and autophagy. Reduction of cellular cholesterol content by simvastatin (green arrows) inhibits mTORC1 activation and induces AMPK activation, both of which lead to increased nuclear translocation of TFEB to induce the expression of autophagy-related genes. *M. tuberculosis* infection (red arrows) induces opposing effects, as it blocks AMPK and induces mTORC1 activation to prevent nuclear translocation of TFEB and block autophagy. For simplicity, the effects of AMPK on mTORC1 activation status, and vice versa, are not shown.

Our findings fully agree with the known mechanistic links between cellular cholesterol homeostasis and mTORC1 signaling. The lysosome is the cellular site for mTORC1 activation (53) as well as a major sorting site for cellular cholesterol through the Nieman-Pick C (NPC) multi-protein machinery (61). Cholesterol homeostasis is critical for activating mTORC1, since depleting cellular cholesterol with methyl-beta-cyclodextrin (62) and blocking cholesterol egress from the lysosome with itraconazole (63) or by siRNA-mediated knockdown of the NPC genes (63) all suppress mTORC1 signaling. We find that statins inhibit mTORC1 signaling at cholesterol-lowering doses and in a cholesterol-dependent manner. Moreover, the compound U18666A, which inhibits the NPC machinery (64), phenocopies the statin effects on autophagy and *M. tuberculosis* burden. Thus, mTORC1 senses cholesterol content and cholesterol trafficking inside the cell, while statins inhibit mTORC1 signaling by dysregulating cholesterol homeostasis. It is worth noting that, while our work and the above-cited literature strongly link mTORC1 inhibition by statins with their cholesterol-lowering activity, a prior report shows that statins inhibit mTORC1 signaling through protein prenylation, at least in coronary arterial myocytes (24). Thus, condition-dependent mechanisms may exist.

Our results fundamentally differ from a previous report proposing the involvement of geranylgeranyl biosynthesis in statin-mediated induction of autophagy during *M. tuberculosis* infection (9). Our conclusions are strongly supported by the findings that the anti-mycobacterial activity of statins is phenocopied only by cholesterol-branch but not prenylation-branch inhibitors and that the biological effects of simvastatin on the AMPK-mTORC1-TFEB-autophagy axis are reversed by adding exogenous cholesterol to the cells. We propose that the different conclusions reached by the previous authors most likely depend on their employment of murine cells and very high simvastatin doses (500-fold higher than the dose we used), which cannot distinguish between anti-cholesterol and anti-prenylation effects of the statin, are toxic for human (THP1) cells, and far exceed the plasma levels attained during therapeutic administration of simvastatin (65).

Our work shows that the cholesterol-lowering activity of statins can affect autophagy in multiple, interconnected ways. First, statins induce AMPK signaling by increasing expression of LKB1 and AMPK genes and the ratio of intracellular AMP:ATP, which activates AMPK (66). The effect on the AMP:ATP ratio is presumably achieved by impairing mitochondrial function, as previously shown (57). Second, statins inhibit mTORC1 activation by decreasing cholesterol biosynthesis and altering homeostasis of cellular cholesterol, as we discussed above. Third, mTORC1 inhibition may also occur by activating AMPK signaling, which prevents mTORC1 activation (67). In addition, a double negative feedback loop between mTORC1 and AMPK has been recently proposed (68). Statins fully counter the effects of *M. tuberculosis* infection on the key AMPK-mTORC1-autophagy axis (Fig. 7), which strongly supports the potential value of this drug class as anti-tubercular therapeutics.

A role for TFEB in the anti-tubercular activity of statins, as demonstrated in the present work, implies that TFEB may be classified as a target for anti-tuberculosis therapeutic intervention. Since TFEB directly regulates autophagy-inducing effector functions, compounds targeting TFEB may exhibit fewer pleiotropic effects – and presumably fewer unwanted side effects -- than those targeting upstream factors like mTORC1. Indeed, everolimus, a rapamycin-analog (69) that is under consideration for use in host-directed therapy against tuberculosis (70), may cause immunosuppression (71) and facilitate reactivation of latent *M. tuberculosis* infection (72). Thus, further investigation will be needed to determine the therapeutic dose of everolimus and the optimal mode of its delivery. The work presented here not only provides the molecular basis for the anti-tubercular activity of statins, but also fosters investigations into novel therapeutic strategies to combat tuberculosis.

The mTORC1-AMPK-TFEB axis has emerged as a key lysosome-based regulator of many biological processes, including autophagy, cellular metabolism, immune responses, inflammation, and damage resolution (73-75). In addition, this axis has a functional impact on host-pathogen responses. For example, AMPK activation is beneficial in infections against hepatitis B and C viruses, but it is detrimental with Dengue and Ebola viruses (76). In another example, *Leishmania major* produces proteases that block mTOR activation, thus repressing the type 1 interferon response and enabling the parasite to survive inside cells (77). *Listeria monocytogenes* and *Staphylococcus aureus* stimulate mTOR by promoting activation of phosphoinositide 3-kinases (PI3K)/protein kinase B (Akt) signaling pathways, which favor pathogen survival during infection (78). The changed perspective on statin-mediated modulation of the mTORC1-AMPK-TFEB axis resulting from the present work will spur novel knowledge-based strategies of host-directed therapies against various infectious agents.

## 4. METHODS

### 4.1. Antibodies and reagents

Simvastatin hydroxy acid was purchased from Santa Cruz Biotechnology (Dallas, TX). GGTI 298 trifluoroacetate salt hydrate and U18666A were purchased from Cayman Chemical Company (Ann Arbor MI). FTI-277 trifluoroacetate salt, BM15766 sulfate, water soluble cholesterol–methyl-β-cyclodextrin and L-arginine were obtained from Sigma-Aldrich (St. Louis, MO). Digoxin, dorsomorphin, and everolimus were purchased from SelleckChem (Houston, TX). Cholesterol quantification kit was purchased from BioVision Inc. (Milpitas, CA) and performed according to the manufacturer’s protocol. Antibodies against mammalian target of rapamycin (mTOR, catalog #2983), phospho-mTOR (pmTOR, Ser2448, catalog #2971), transcription factor EB (tfeb, catalog #32361), sequestosome 1 (SQSTM1/p62, catalog #88588), beta actin (β-actin, catalog #4970) and lamin B1 (catalog #12586) were purchased from Cell Signaling Technology (Danvers, MA). Antibodies against Rab escort protein 1 (REP-1, catalog #sc-23905), Ras homolog family member B (RhoB, catalog #sc-180), Ras-related protein Rap-1A (catalog #sc-14872-R, specific for unprenylated Rap1a) and Ras-related protein Rab5 (catalog #sc-46692), were purchased from Santa Cruz Biotechnology (Dallas TX). Secondary antibodies IRDye® 800CW goat anti-rabbit IgG, IRDye® 680RD donkey anti-mouse IgG and IRDye® 800CW goat anti-mouse IgG were purchased from LI-COR Biosciences (Lincoln, NE).

### 4.2. Culturing of cell lines

THP1 monocytes were purchased from American Type Culture Collection (ATCC, Manassas, VA). THPI monocytes were grown in RPMI 1640 culture medium (Corning, Manassas, VA) supplemented with 4 mM L-glutamine (Corning, Manassas, VA), 10% FBS (Seradigm, Radnor, PA) and penicillin-streptomycin solution (Corning, Manassas, VA) incubated at 37°C in a humidified atmosphere consisting of 5% CO2. For differentiation, phorbol-12-myristate 13-acetate (PMA) (Sigma-Aldrich, St. Louis, MO) was added overnight at a final concentration of 40 nM.

### 4.3. siRNA transfection

Differentiated THP1 cells, seeded at a density of 3.3 × 10^5^ cells/well in 6-well plates were in RPMI-1640 supplemented with 2% fetal bovine serum (Seradigm, Radnor, PA)and 4 mM L-glutamine (Corning, Manassas, VA). Commercially available ON-TARGET plus Non-targeting and CHM SMARTpool siRNA (Dharmacon, Lafayette, CO) was reconstituted with 1X siRNA buffer (Dharmacon, Lafayette, CO) to a stock concentration of 20µM. Transfection with siRNA was performed at a final concentration of 50 nM using DharmaFECT2 Transfection reagent (Dharmacon, Lafayette, CO) according to the manufacturer’s protocol. After 72 hours, cells were subjected to a second round of transfection with an identical concentration (50 nM) of siRNA. The second round of siRNA treatment was performed to prolong the duration of silencing the target protein at day 6 of sample collection. After 6 days, the cells were assayed for protein expression by Western blot analysis, as described below.

### 4.4. *In vitro* infection of differentiated THP1

Differentiated THP1 cells, seeded at a density of 5 × 10^5^ cells/well in 24-well plates were infected with a frozen stock of *M. tuberculosis* H37Rv at a multiplicity of infection (MOI) of 1:20, as previously described (11, 79). The bacterial inoculum for infection was prepared by diluting a frozen bacterial stock in supplemented RPMI-1640 medium (as above) to obtain the desired MOI. Bacterial clumps were disrupted by vortexing with sterile 3-mm-diameter glass beads for 2 min. The resulting suspension was used for infection of primary and culture cells. Infected cells were washed three times, at 4 hours post-infection, to remove extracellular bacteria and were incubated with fresh RPMI-1640 supplemented with 2% fetal bovine serum (Seradigm, Radnor, PA) and 4 mM L-glutamine (Corning, Manassas, VA) and treated with either chemical inhibitors or solvent controls. Media was replenished at days 3 post infection. At day 6 post-infection, cells were lysed with 0.05% SDS in water and serial dilutions plated on 7H10 agar plates. After three weeks, colony-forming units (CFU) were enumerated. Cell viability and total number of cells were determined using trypan blue counting.

### 4.5. Preparation of whole cell lysate and subcellular fractions

THP1 cells seeded in 6-well plates at a density of 2 × 10^6^ cells/well were treated with compounds for the duration of the experiment. A total of 2 × 10^6^ macrophages were lysed in 100μL RIPA lysis buffer (Santa Cruz Biotechnology, Dallas, TX). Subcellular Protein Fractionation Kit for Cultured Cells (Thermo Fischer Scientific, Waltham, MA) was used to prepare the various subcellular protein extracts according to the manufacturer’s instructions. Protein lysates were filtered and sterilized through a 0.2 μm filter and stored at −20 °C.

### 4.6. Western blot analysis

We used the BCA Protein Assay (Thermo Fischer Scientific, Waltham, MA) to determine the protein concentration. Equal amounts of protein were then resolved by SDS-PAGE on either 7.5% or 10% polyacrylamide gels (Bio-Rad, Hercules, CA). Separated proteins were transferred to PVDF Immobilon-FL membranes (Millipore, Billerica, MA) using the BioRad Trans-Blot Turbo Transfer System (Hercules, CA). The membrane was blocked with Odyssey Block buffer (LI-COR Biosciences, Lincoln, NE) at RT for 2 hours and incubated with primary antibodies (1:1000) at 4°C overnight. The membrane was then incubated with secondary near-infrared fluorescent antibodies (1:10,000) at room temperature for 1 hour. Immunoblots were scanned using a LI-COR Odyssey Infrared Imager (Lincoln, NE). Antibody signals were quantified using LI-COR Odyssey Infrared Imager software version 1.2. Quantitated protein levels were normalized to total protein levels using REVERT total protein stain (LI-COR Biosciences, Lincoln, NE), beta-actin or lamin B1. Immunoblots were stripped with NewBlot PVDF stripping buffer (Lincoln, NE) and reprobed when required.

### 4.7. AMP/ATP assay

Cellular AMP and ATP was extracted using the boiling water method (80). Cells were seeded onto 12-well plates and treated with simvastatin for 6 days. Cells were washed twice with cold PBS, followed by the addition of ice-cold water. Cells were scraped and collected into pre-chilled tube and were lysed by vigorous vortexing on ice for 10 seconds. The protein concentration was determined by the BCA Protein Assay (Thermo Fischer Scientific, Waltham, MA) using 10 μL of the lysate. The remaining lysate was boiled with shaking for 10 minutes, cooled on ice for 30 seconds and centrifuged at 13000 rpm for 5 minutes. The supernatant was collected and stored at −80°C until further use. The levels of ATP, ADP and AMP were determined using an ATP/ADP/AMP Assay Kit (Biomedical Research Service Center, University at Buffalo, State University of New York). The luciferase bioluminescence was measured using the BioTek Synergy H1 microplate reader and was compared with a standard curve of ATP concentrations. AMP concentration was calculated according to the manufacturer’s instruction.

### 4.8. Statistical analysis

All values are presented as means ± standard deviation (SD) (*n*=3). Comparisons between two groups were performed using a two-tailed Student’s *t*-test. The criterion for statistical significance was *P* < 0.05.

### 4.9. Transcriptome data analysis

Statistical analysis was performed in R (*) using the Knowledge Synthesis KS/prot data integration and discovery platform (Knowledge Synthesis Inc., Berkeley, CA, USA; knowledgesynthesis.com). T-tests were performed to compare measurements of transcript levels between sample groups, and the resulting p-values were calculated. To explore the association between simvastatin and specific protein functions and pathways, an experimental compendium of gene-set analyses of tuberculosis-relevant experiments, including simvastatin-treated macrophages, was established in a searchable results repository, iDataMed/tb (Knowledge Synthesis Inc., Berkeley, CA, USA; knowledgesynthesis.com). The compendium was comprised of results calculated in the same manner as in previous research (79, 81, 82) and included analyses of different types of gene sets defined by NCBI biosystems (https://www.ncbi.nlm.nih.gov/biosystems), Reactome (http://reactome.org/), and transcriptional modulators (83). Each gene set was tested for extreme ranks of differential expression among all measured genes in each comparison by coincident extreme ranks in numerical observations (CERNO). Multiple transcript measurements were combined as described previously (82, 84). The Benjamini–Hochberg method was used to calculate the false discovery rate (FDR) within each gene set type. Search was facilitated in iDataMed/tb by a text query interface, in which searches such as simvastatin AND mTOR or simvastatin AND ribosome biogenesis returned statistically significant findings sorted in a fashion weighted to present top-ranking and more significant results first.

## Acknowledgments

We thank Shumin Tan for critical reading of the manuscript and Life Science Editors for editing assistance. This work was funded by NIH grants to PK (UH2/3 AI122309) and MLG (R01 HL149450; R01AI104615; UL1TR003017) and a New Jersey Health Foundation grant to MLG (PC 10-15).

---

**Supplementary Figure 1.**
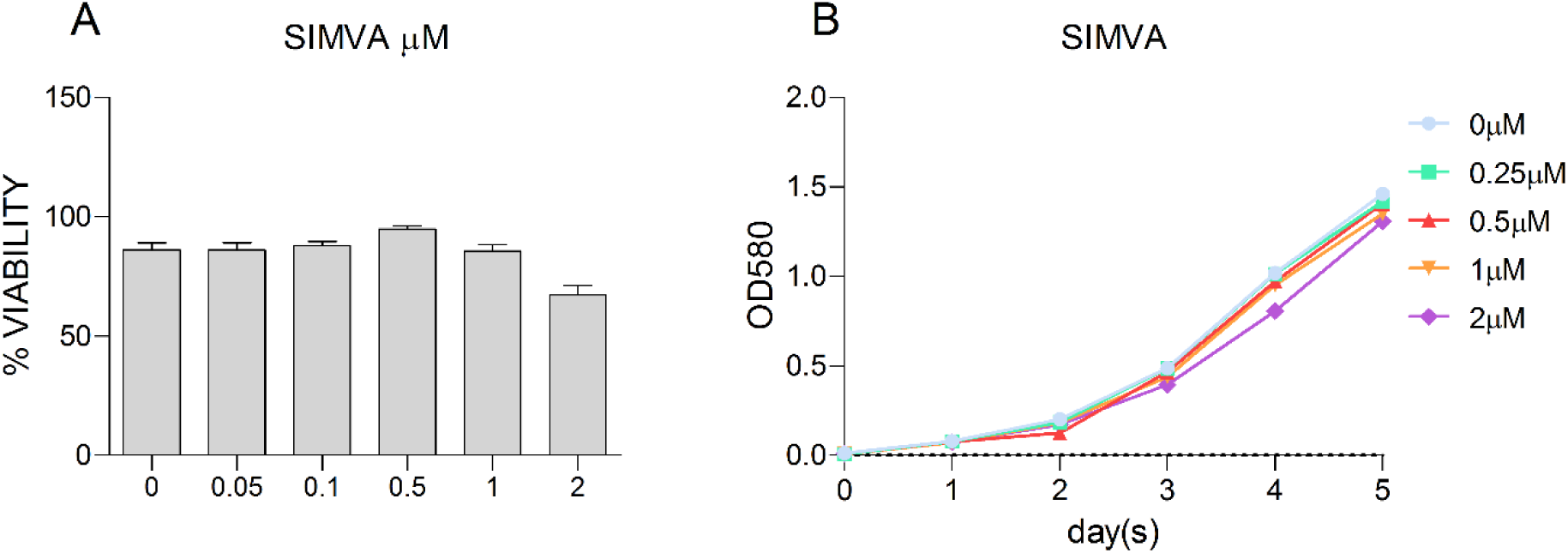
Effect of simvastatin on *M. tuberculosis* axenic cultures and toxicity of THP1 cells. (**A**) *M. tuberculosis* at an optical density (OD) 580 of 0.01 was exposed to simvastatin at the indicated doses for 5 days. Growth of the bacilli was recorded by using a spectrophotometer. (**B**) Cell viability of infected THP1 cells treated with simvastatin at the indicated doses for 6 days calculated was evaluated using trypan blue exclusion (viability was affected at 2µM).

**Supplementary Figure 2.**
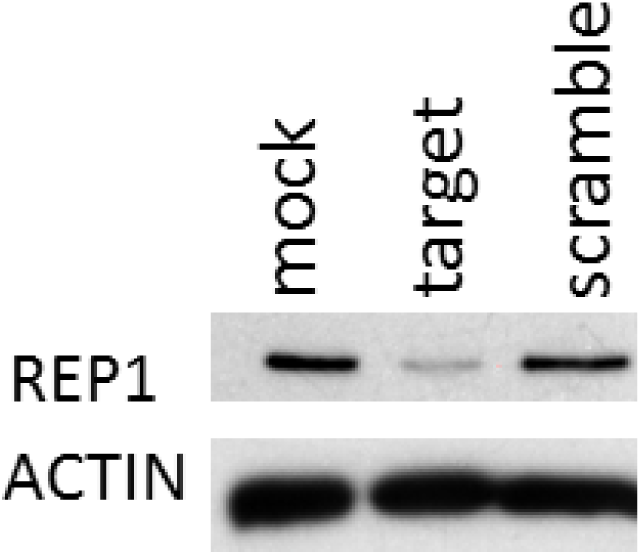
siRNA-mediated protein knockdown. Western blot analysis of REP-1 and beta-actin. Lane 1: mock transfected THP1 cells with transfection reagent, lane 2: THP1 cells transfected with targeting siRNA, and lane 3: THP1 cells transfected with non-targeting siRNA.

**Supplementary Table 1.** Calculated p-values were determined using the one-sided T-test to compare transcript levels between simvastatin treatment and solvent control. P-values for multiple transcripts for each gene measured on the microarray platform were first integrated as in a nested CERNO test and sorted as described in Methods.

| Gene | GeneID | MinProbeP | MinProbeP_BH |
| --- | --- | --- | --- |
| GRB10 | 2887 | 0.001109824 | 0.430328378 |
| ULK2 | 9706 | 0.000932144 | 0.421150989 |
| SESN2 | 83667 | 0.001436966 | 0.443984358 |
| IGF1R | 3480 | 0.004732249 | 0.450956278 |
| EIF4EBP1 | 1978 | 0.001242791 | 0.443984358 |
| INSR | 3643 | 0.013018491 | 0.47852084 |
| BRAF | 673 | 0.001555621 | 0.443984358 |
| SLC7A5 | 8140 | 0.002160195 | 0.443984358 |
| RPS6 | 6194 | 0.017128786 | 0.483459095 |
| PTEN | 5728 | 0.011421643 | 0.47852084 |
| FZD7 | 8324 | 0.015526197 | 0.482266701 |
| KRAS | 3845 | 0.001753226 | 0.443984358 |
| ULK1 | 8408 | 0.006171159 | 0.469816424 |
| MLST8 | 64223 | 0.00629089 | 0.469816424 |
| AKT3 | 10000 | 0.001886011 | 0.443984358 |
| PRKCB | 5579 | 0.025957045 | 0.497540389 |
| RPS6KA2 | 6196 | 0.026284887 | 0.498394638 |
| EIF4E2 | 9470 | 0.013837528 | 0.47852084 |
| MAPKAP1 | 79109 | 0.025432697 | 0.495000508 |
| RRAGD | 58528 | 0.023325731 | 0.491452839 |
| AKT2 | 208 | 0.009170944 | 0.469816424 |
| STRADB | 55437 | 0.017106803 | 0.483459095 |
| DDIT4 | 54541 | 0.017529493 | 0.484205568 |
| RRAGB | 10325 | 0.018359747 | 0.485255556 |
| SLC38A9 | 153129 | 0.034598298 | 0.51335088 |
| TNFRSF1A | 7132 | 0.019111025 | 0.485255556 |
| MAPK3 | 5595 | 0.021819564 | 0.489762465 |
| PIK3R1 | 5295 | 0.0407176 | 0.522684349 |
| RRAGC | 64121 | 0.048074372 | 0.533371235 |
| LAMTOR1 | 55004 | 0.033429789 | 0.512020729 |
| SOS2 | 6655 | 0.030070733 | 0.506764922 |
| DEPTOR | 64798 | 0.012097783 | 0.47852084 |
| EIF4B | 1975 | 0.033837131 | 0.5122758 |
| LAMTOR2 | 28956 | 0.060804675 | 0.55276242 |
| MAP2K2 | 5605 | 0.074739787 | 0.574982329 |
| NPRL2 | 10641 | 0.043228173 | 0.524298087 |
| WNT6 | 7475 | 0.031691656 | 0.511284177 |
| WDR59 | 79726 | 0.028230583 | 0.503194159 |
| FZD1 | 8321 | 0.075315677 | 0.5755359 |
| DVL2 | 1856 | 0.088263221 | 0.581068377 |
| WNT2B | 7482 | 0.021182142 | 0.488465265 |
| ATP6V1E2 | 90423 | 0.11925257 | 0.593878314 |
| TSC2 | 7249 | 0.127442843 | 0.599279544 |
| DEPDC5 | 9681 | 0.137480168 | 0.610375159 |
| FNIP1 | 96459 | 0.01838442 | 0.485255556 |
| WNT5A | 7474 | 0.051121338 | 0.538393653 |
| WNT10A | 80326 | 0.117448245 | 0.593115239 |
| WNT8B | 7479 | 0.15086932 | 0.624017139 |
| RPS6KA6 | 27330 | 0.070306664 | 0.568531923 |
| RNF152 | 220441 | 0.080750723 | 0.581068377 |
| LAMTOR3 | 8649 | 0.153425297 | 0.626745609 |
| FZD5 | 7855 | 0.06031968 | 0.551298914 |
| SOS1 | 6654 | 0.049490311 | 0.535249151 |
| NRAS | 4893 | 0.097327924 | 0.581068377 |
| SLC3A2 | 6520 | 0.189841661 | 0.662664415 |
| MAP2K1 | 5604 | 0.191960298 | 0.664760673 |
| WDR24 | 84219 | 0.203396803 | 0.676407272 |
| WNT9B | 7484 | 0.210123965 | 0.67906154 |
| IRS1 | 3667 | 0.096786186 | 0.581068377 |
| RPS6KA1 | 6195 | 0.214463557 | 0.681974213 |
| FZD2 | 2535 | 0.165364898 | 0.638522599 |
| PIK3R2 | 5296 | 0.100054089 | 0.581386593 |
| ATP6V1C2 | 245973 | 0.148829059 | 0.621413037 |
| AKT1S1 | 84335 | 0.20429009 | 0.677398581 |
| WNT1 | 7471 | 0.274083847 | 0.7378545 |
| NPRL3 | 8131 | 0.16724259 | 0.640306024 |
| FLCN | 201163 | 0.150405227 | 0.623509226 |
| TBC1D7 | 51256 | 0.159742801 | 0.632967078 |
| FZD4 | 8322 | 0.12268478 | 0.595148053 |
| PRKCG | 5582 | 0.175996141 | 0.649562782 |
| WNT4 | 54361 | 0.118067005 | 0.593115239 |
| WNT2 | 7472 | 0.320296942 | 0.776474926 |
| FZD10 | 11211 | 0.325652817 | 0.780762987 |
| CHUK | 1147 | 0.338654657 | 0.791085931 |
| RHEB | 6009 | 0.079515595 | 0.581068377 |
| PIK3CA | 5290 | 0.072933318 | 0.572924748 |
| WNT5B | 81029 | 0.120101513 | 0.59429342 |
| GSK3B | 2932 | 0.070470671 | 0.568595442 |
| WNT7A | 7476 | 0.392161441 | 0.831620814 |
| RHOA | 387 | 0.188268426 | 0.66052823 |
| SEH1L | 81929 | 0.281114595 | 0.74463589 |
| CLIP1 | 6249 | 0.069498256 | 0.568531923 |
| ATP6V1C1 | 528 | 0.072178463 | 0.570914588 |
| RRAGA | 10670 | 0.440891437 | 0.867013879 |
| ATP6V1D | 51382 | 0.099906682 | 0.581386593 |
| RPS6KB2 | 6199 | 0.476173439 | 0.889098517 |
| INS | 3630 | 0.477696635 | 0.889894726 |
| LRP5 | 4041 | 0.278231083 | 0.741804826 |
| MAPK1 | 5594 | 0.072083034 | 0.570555939 |
| WNT11 | 7481 | 0.532316263 | 0.919227738 |
| ATP6V1B2 | 526 | 0.536598735 | 0.92153058 |
| SGK1 | 6446 | 0.542085215 | 0.923699015 |
| WNT8A | 7478 | 0.544991102 | 0.925230986 |
| DVL3 | 1857 | 0.291089352 | 0.752366244 |
| AKT1 | 207 | 0.552424577 | 0.928954124 |
| WNT10B | 7480 | 0.569621293 | 0.9383551 |
| FZD9 | 8326 | 0.583207883 | 0.94466096 |
| ATP6V1B1 | 525 | 0.290341236 | 0.751854121 |
| PDPK1 | 5170 | 0.611189618 | 0.954444297 |
| IKBKB | 3551 | 0.289651865 | 0.751029291 |
| TSC1 | 7248 | 0.623227067 | 0.960129794 |
| PIK3CB | 5291 | 0.373978832 | 0.818307275 |
| STK11 | 6794 | 0.199244725 | 0.672372544 |
| WNT16 | 51384 | 0.418934473 | 0.851750995 |
| PRKCA | 5578 | 0.129900376 | 0.601143843 |
| GRB2 | 2885 | 0.125690656 | 0.598327418 |
| HRAS | 3265 | 0.695605065 | 0.986581513 |
| MTOR | 2475 | 0.354796062 | 0.803898789 |
| SKP2 | 6502 | 0.243846648 | 0.711144578 |
| ATP6V1G3 | 127124 | 0.719031911 | 0.993979891 |
| ATP6V1G2 | 534 | 0.730911414 | 0.997595288 |
| RAF1 | 5894 | 0.305262064 | 0.764579102 |
| WNT7B | 7477 | 0.365826655 | 0.811839376 |
| CAB39 | 51719 | 0.518241681 | 0.911390542 |
| LRP6 | 4040 | 0.410719583 | 0.845363548 |
| DVL1 | 1855 | 0.822491981 | 0.999996506 |
| SEC13 | 6396 | 0.832058292 | 0.999996506 |
| LAMTOR4 | 389541 | 0.839767875 | 0.999996506 |
| RPS6KA3 | 6197 | 0.561248385 | 0.933500843 |
| RPS6KB1 | 6198 | 0.463941937 | 0.881512253 |
| CAB39L | 81617 | 0.23092078 | 0.698891595 |
| LAMTOR5 | 10542 | 0.699713516 | 0.988252244 |
| IGF1 | 3479 | 0.29278256 | 0.753931023 |
| TELO2 | 9894 | 0.725863069 | 0.996436457 |
| WNT9A | 7483 | 0.709192336 | 0.990926278 |
| ATP6V1E1 | 529 | 0.942185572 | 0.999996506 |
| ATP6V1G1 | 9550 | 0.811289889 | 0.999996506 |
| RPTOR | 57521 | 0.955653562 | 0.999996506 |
| TTI1 | 9675 | 0.961885455 | 0.999996506 |
| MIOS | 54468 | 0.346511001 | 0.797425998 |
| EIF4E | 1977 | 0.341168316 | 0.793392859 |
| STRADA | 92335 | 0.67751961 | 0.978906917 |
| PRKAA2 | 5563 | 0.377930069 | 0.820872939 |
| ATP6V1H | 51606 | 0.455557934 | 0.876178968 |
| ATP6V1A | 523 | 0.847559077 | 0.999996506 |
| FZD6 | 8323 | 0.981719961 | 0.999996506 |
| FZD3 | 7976 | 0.592488057 | 0.948091877 |
| RICTOR | 253260 | 0.473910736 | 0.887761008 |
| PIK3CD | 5293 | 0.831190851 | 0.999996506 |
| PIK3R3 | 8503 | 0.907976527 | 0.999996506 |
| PRR5 | 55615 | 0.918227678 | 0.999996506 |
| TNF | 7124 | 0.996733204 | 0.999996506 |
| FNIP2 | 57600 | 0.576956264 | 0.941554496 |
| PRKAA1 | 5562 | 0.471421016 | 0.886175554 |
| LPIN1 | 23175 | 0.957125372 | 0.999996506 |

## References

1. Philips JA, Ernst JD. Tuberculosis pathogenesis and immunity. Annu Rev Pathol. 2012;7:353-84, 10.1146/annurev-pathol-011811-132458.

2. Cadena AM, Fortune SM, Flynn JL. Heterogeneity in tuberculosis. Nat Rev Immunol. 2017;17(11):691-702, 10.1038/nri.2017.69.

3. Karakousis PC, Yoshimatsu T, Lamichhane G, Woolwine SC, Nuermberger EL, Grosset J, Bishai WR. Dormancy phenotype displayed by extracellular Mycobacterium tuberculosis within artificial granulomas in mice. J Exp Med. 2004;200(5):647-57, 10.1084/jem.20040646.

4. Lin PL, Ford CB, Coleman MT, Myers AJ, Gawande R, Ioerger T, Sacchettini J, Fortune SM, Flynn JL. Sterilization of granulomas is common in active and latent tuberculosis despite within-host variability in bacterial killing. Nat Med. 2014;20(1):75-9, 10.1038/nm.3412.

5. Gengenbacher M, Kaufmann SHE. Mycobacterium tuberculosis: success through dormancy. FEMS Microbiol Rev. 2012;36(3):514-32, 10.1111/j.1574-6976.2012.00331.x.

6. Mitchison D, Davies G. The chemotherapy of tuberculosis: past, present and future. Int J Tuberc Lung Dis. 2012;16(6):724-32, 10.5588/ijtld.12.0083.

7. World Health O. Global tuberculosis report 2018. Geneva: World Health Organization; 2018 2018.

8. Tiberi S, du Plessis N, Walzl G, Vjecha MJ, Rao M, Ntoumi F, Mfinanga S, Kapata N, Mwaba P, McHugh TD, Ippolito G, Migliori GB, Maeurer MJ, Zumla A. Tuberculosis: progress and advances in development of new drugs, treatment regimens, and host-directed therapies. Lancet Infect Dis. 2018;18(7):e183-e98, 10.1016/S1473-3099(18)30110-5.

9. Parihar SP, Guler R, Khutlang R, Lang DM, Hurdayal R, Mhlanga MM, Suzuki H, Marais AD, Brombacher F. Statin therapy reduces the mycobacterium tuberculosis burden in human macrophages and in mice by enhancing autophagy and phagosome maturation. J Infect Dis. 2014;209(5):754-63, 10.1093/infdis/jit550.

10. Skerry C, Pinn ML, Bruiners N, Pine R, Gennaro ML, Karakousis PC. Simvastatin increases the in vivo activity of the first-line tuberculosis regimen. J Antimicrob Chemother. 2014;69(9):2453-7, 10.1093/jac/dku166.

11. Dutta NK, Bruiners N, Pinn ML, Zimmerman MD, Prideaux B, Dartois V, Gennaro ML, Karakousis PC. Statin adjunctive therapy shortens the duration of TB treatment in mice. J Antimicrob Chemother. 2016;71(6):1570-7, 10.1093/jac/dkw014.

12. Dutta NK, Bruiners N, Zimmerman MD, Tan S, Dartois V, Gennaro ML, Karakousis PC. Adjunctive host-directed therapy with statins improves tuberculosis-related outcomes in mice. J Infect Dis. 2019:jiz517. 10.1093/infdis/jiz517.

13. Guerra-De-Blas PDC, Bobadilla-Del-Valle M, Sada-Ovalle I, Estrada-García I, Torres-González P, López-Saavedra A, Guzmán-Beltrán S, Ponce-de-León A, Sifuentes-Osornio J. Simvastatin Enhances the Immune Response Against Mycobacterium tuberculosis. Front Microbiol. 2019;10:2097-, 10.3389/fmicb.2019.02097.

14. Willey JZ, Elkind MSV. 3-Hydroxy-3-methylglutaryl-coenzyme A reductase inhibitors in the treatment of central nervous system diseases. Arch Neurol. 2010;67(9):1062-7, 10.1001/archneurol.2010.199.

15. Xu N, Shen N, Wang X, Jiang S, Xue B, Li C. Protein prenylation and human diseases: a balance of protein farnesylation and geranylgeranylation. Sci China Life Sci. 2015;58(4):328-35, 10.1007/s11427-015-4836-1.

16. Zhou Q, Liao JK. Pleiotropic effects of statins. - Basic research and clinical perspectives. Circ J. 2010;74(5):818-26, 10.1253/circj.cj-10-0110.

17. Blanco-Colio LM, Tuñón J, Martín-Ventura JL, Egido J. Anti-inflammatory and immunomodulatory effects of statins. Kidney Int. 2003;63(1):12-23, 10.1046/j.1523-1755.2003.00744.x.

18. Rachoin J-S, Cerceo E, Dellinger RP. A new role for statins in sepsis. Crit Care. 2013;17(1):105-, 10.1186/cc11907.

19. Yende S, Milbrandt EB, Kellum JA, Kong L, Delude RL, Weissfeld LA, Angus DC. Understanding the potential role of statins in pneumonia and sepsis. Crit Care Med. 2011;39(8):1871-8, 10.1097/CCM.0b013e31821b8290.

20. Zeiser R. Immune modulatory effects of statins. Immunology. 2018;154(1):69-75, 10.1111/imm.12902.

21. Castillo EF, Dekonenko A, Arko-Mensah J, Mandell MA, Dupont N, Jiang S, Delgado-Vargas M, Timmins GS, Bhattacharya D, Yang H, Hutt J, Lyons CR, Dobos KM, Deretic V. Autophagy protects against active tuberculosis by suppressing bacterial burden and inflammation. Proc Natl Acad Sci U S A. 2012;109(46):E3168-E76, 10.1073/pnas.1210500109.

22. Bento CF, Empadinhas N, Mendes V. Autophagy in the fight against tuberculosis. DNA Cell Biol. 2015;34(4):228-42, 10.1089/dna.2014.2745.

23. Vergne I, Chua J, Lee H-H, Lucas M, Belisle J, Deretic V. Mechanism of phagolysosome biogenesis block by viable Mycobacterium tuberculosis. Proc Natl Acad Sci U S A. 2005;102(11):4033-8, 10.1073/pnas.0409716102.

24. Wei Y-M, Li X, Xu M, Abais JM, Chen Y, Riebling CR, Boini KM, Li P-L, Zhang Y. Enhancement of autophagy by simvastatin through inhibition of Rac1-mTOR signaling pathway in coronary arterial myocytes. Cell Physiol Biochem. 2013;31(6):925-37, 10.1159/000350111.

25. Parikh A, Childress C, Deitrick K, Lin Q, Rukstalis D, Yang W. Statin-induced autophagy by inhibition of geranylgeranyl biosynthesis in prostate cancer PC3 cells. Prostate. 2010;70(9):971-81, 10.1002/pros.21131.

26. Bloch H, Segal W. Biochemical differentiation of Mycobacterium tuberculosis grown in vivo and in vitro. J Bacteriol. 1956;72(2):132–41,

27. Nazarova EV, Montague CR, La T, Wilburn KM, Sukumar N, Lee W, Caldwell S, Russell DG, VanderVen BC. Rv3723/LucA coordinates fatty acid and cholesterol uptake in Mycobacterium tuberculosis. Elife. 2017;6:e26969. 10.7554/eLife.26969.

28. Wipperman MF, Sampson NS, Thomas ST. Pathogen roid rage: cholesterol utilization by Mycobacterium tuberculosis. Crit Rev Biochem Mol Biol. 2014;49(4):269-93, 10.3109/10409238.2014.895700.

29. Griffin JE, Pandey AK, Gilmore SA, Mizrahi V, McKinney JD, Bertozzi CR, Sassetti CM. Cholesterol catabolism by Mycobacterium tuberculosis requires transcriptional and metabolic adaptations. Chem Biol. 2012;19(2):218-27, 10.1016/j.chembiol.2011.12.016.

30. Pandey AK, Sassetti CM. Mycobacterial persistence requires the utilization of host cholesterol. Proc Natl Acad Sci U S A. 2008;105(11):4376-80, 10.1073/pnas.0711159105.

31. Miner MD, Chang JC, Pandey AK, Sassetti CM, Sherman DR. Role of cholesterol in Mycobacterium tuberculosis infection. Indian J Exp Biol. 2009;47(6):407–11,

32. Masiewicz P, Brzostek A, Wolański M, Dziadek J, Zakrzewska-Czerwińska J. A novel role of the PrpR as a transcription factor involved in the regulation of methylcitrate pathway in Mycobacterium tuberculosis. PloS one. 2012;7(8):e43651-e, 10.1371/journal.pone.0043651.

33. Datta P, Shi L, Bibi N, Balázsi G, Gennaro ML. Regulation of central metabolism genes of Mycobacterium tuberculosis by parallel feed-forward loops controlled by sigma factor E (σ(E)). J Bacteriol. 2011;193(5):1154-60, 10.1128/JB.00459-10.

34. Paik S, Kim JK, Chung C, Jo E-K. Autophagy: A new strategy for host-directed therapy of tuberculosis. Virulence. 2019;10(1):448-59, 10.1080/21505594.2018.1536598.

35. Bjørkøy G, Lamark T, Pankiv S, Øvervatn A, Brech A, Johansen T. Monitoring autophagic degradation of p62/SQSTM1. Methods Enzymol. 2009;452:181-97, 10.1016/S0076-6879(08)03612-4.

36. Dong H, Jing W, Runpeng Z, Xuewei X, Min M, Ru C, Yingru X, Shengfa N, Rongbo Z. ESAT6 inhibits autophagy flux and promotes BCG proliferation through MTOR. Biochem Biophys Res Commun. 2016;477(2):195-201, 10.1016/j.bbrc.2016.06.042.

37. Sparrow SM, Carter JM, Ridgway ND, Cook HW, Byers DM. U18666A inhibits intracellular cholesterol transport and neurotransmitter release in human neuroblastoma cells. Neurochem Res. 1999;24(1):69-77, 10.1023/a:1020932130753.

38. Heitman J, Movva NR, Hall MN. Targets for cell cycle arrest by the immunosuppressant rapamycin in yeast. Science. 1991;253(5022):905-9, 10.1126/science.1715094.

39. Choe G, Horvath S, Cloughesy TF, Crosby K, Seligson D, Palotie A, Inge L, Smith BL, Sawyers CL, Mischel PS. Analysis of the phosphatidylinositol 3’-kinase signaling pathway in glioblastoma patients in vivo. Cancer Res. 2003;63(11):2742–6,

40. Zhou X, Tan M, Stone Hawthorne V, Klos KS, Lan K-H, Yang Y, Yang W, Smith TL, Shi D, Yu D. Activation of the Akt/mammalian target of rapamycin/4E-BP1 pathway by ErbB2 overexpression predicts tumor progression in breast cancers. Clin Cancer Res. 2004;10(20):6779-88, 10.1158/1078-0432.CCR-04-0112.

41. Kong X, Tan B, Yin Y, Gao H, Li X, Jaeger LA, Bazer FW, Wu G. L-Arginine stimulates the mTOR signaling pathway and protein synthesis in porcine trophectoderm cells. J Nutr Biochem. 2012;23(9):1178-83, 10.1016/j.jnutbio.2011.06.012.

42. Leung EY, Askarian-Amiri M, Finlay GJ, Rewcastle GW, Baguley BC. Potentiation of Growth Inhibitory Responses of the mTOR Inhibitor Everolimus by Dual mTORC1/2 Inhibitors in Cultured Breast Cancer Cell Lines. PloS one. 2015;10(7):e0131400-e, 10.1371/journal.pone.0131400.

43. Kawata T, Tada K, Kobayashi M, Sakamoto T, Takiuchi Y, Iwai F, Sakurada M, Hishizawa M, Shirakawa K, Shindo K, Sato H, Takaori-Kondo A. Dual inhibition of the mTORC1 and mTORC2 signaling pathways is a promising therapeutic target for adult T-cell leukemia. Cancer Sci. 2018;109(1):103-11, 10.1111/cas.13431.

44. Medina DL, Di Paola S, Peluso I, Armani A, De Stefani D, Venditti R, Montefusco S, Scotto-Rosato A, Prezioso C, Forrester A, Settembre C, Wang W, Gao Q, Xu H, Sandri M, Rizzuto R, De Matteis MA, Ballabio A. Lysosomal calcium signalling regulates autophagy through calcineurin and TFEB. Nat Cell Biol. 2015;17(3):288-99, 10.1038/ncb3114.

45. Zhang J, Wang J, Xu J, Lu Y, Jiang J, Wang L, Shen H-M, Xia D. Curcumin targets the TFEB-lysosome pathway for induction of autophagy. Oncotarget. 2016;7(46):75659-71, 10.18632/oncotarget.12318.

46. Napolitano G, Esposito A, Choi H, Matarese M, Benedetti V, Di Malta C, Monfregola J, Medina DL, Lippincott-Schwartz J, Ballabio A. mTOR-dependent phosphorylation controls TFEB nuclear export. Nat Commun. 2018;9(1):3312-, 10.1038/s41467-018-05862-6.

47. Zhitomirsky B, Yunaev A, Kreiserman R, Kaplan A, Stark M, Assaraf YG. Lysosomotropic drugs activate TFEB via lysosomal membrane fluidization and consequent inhibition of mTORC1 activity. Cell Death Dis. 2018;9(12):1191-, 10.1038/s41419-018-1227-0.

48. Tuomisto TT, Lumivuori H, Kansanen E, Häkkinen S-K, Turunen MP, van Thienen JV, Horrevoets AJ, Levonen A-L, Ylä-Herttuala S. Simvastatin has an anti-inflammatory effect on macrophages via upregulation of an atheroprotective transcription factor, Kruppel-like factor 2. Cardiovasc Res. 2008;78(1):175-84, 10.1093/cvr/cvn007.

49. Liu B, Liu F. Feedback regulation of mTORC1 by Grb10 in metabolism and beyond. Cell Cycle. 2014;13(17):2643-4, 10.4161/15384101.2014.954221.

50. Mlitz V, Gendronneau G, Berlin I, Buchberger M, Eckhart L, Tschachler E. The Expression of the Endogenous mTORC1 Inhibitor Sestrin 2 Is Induced by UVB and Balanced with the Expression Level of Sestrin 1. PloS one. 2016;11(11):e0166832-e, 10.1371/journal.pone.0166832.

51. Cheng H, Yang Z-T, Bai Y-Q, Cai Y-F, Zhao J-P. Overexpression of Ulk2 inhibits proliferation and enhances chemosensitivity to cisplatin in non-small cell lung cancer. Oncol Lett. 2019;17(1):79-86, 10.3892/ol.2018.9604.

52. Jung CH, Ro S-H, Cao J, Otto NM, Kim D-H. mTOR regulation of autophagy. FEBS Lett. 2010;584(7):1287-95, 10.1016/j.febslet.2010.01.017.

53. Condon KJ, Sabatini DM. Nutrient regulation of mTORC1 at a glance. J Cell Sci. 2019;132(21):jcs222570. 10.1242/jcs.222570.

54. Vilar E, Perez-Garcia J, Tabernero J. Pushing the envelope in the mTOR pathway: the second generation of inhibitors. Mol Cancer Ther. 2011;10(3):395-403, 10.1158/1535-7163.MCT-10-0905.

55. Yan Q, Han C, Wang G, Waddington JL, Zheng L, Zhen X. Activation of AMPK/mTORC1-Mediated Autophagy by Metformin Reverses Clk1 Deficiency-Sensitized Dopaminergic Neuronal Death. Mol Pharmacol. 2017;92(6):640-52, 10.1124/mol.117.109512.

56. Shackelford DB, Shaw RJ. The LKB1-AMPK pathway: metabolism and growth control in tumour suppression. Nat Rev Cancer. 2009;9(8):563-75, 10.1038/nrc2676.

57. Ma L, Niknejad N, Gorn-Hondermann I, Dayekh K, Dimitroulakos J. Lovastatin induces multiple stress pathways including LKB1/AMPK activation that regulate its cytotoxic effects in squamous cell carcinoma cells. PloS one. 2012;7(9):e46055-e, 10.1371/journal.pone.0046055.

58. Liu X, Chhipa RR, Nakano I, Dasgupta B. The AMPK inhibitor compound C is a potent AMPK-independent antiglioma agent. Mol Cancer Ther. 2014;13(3):596-605, 10.1158/1535-7163.MCT-13-0579.

59. Göransson O, McBride A, Hawley SA, Ross FA, Shpiro N, Foretz M, Viollet B, Hardie DG, Sakamoto K. Mechanism of action of A-769662, a valuable tool for activation of AMP-activated protein kinase. J Biol Chem. 2007;282(45):32549-60, 10.1074/jbc.M706536200.

60. Allergy NIo, Diseases I. StAT-TB (Statin Adjunctive Therapy for TB): A Phase 2b Dose-finding Study of Pravastatin in Adults With Tuberculosis. https://ClinicalTrials.gov/show/NCT03882177; 2020.

61. Sleat DE, Wiseman JA, El-Banna M, Price SM, Verot L, Shen MM, Tint GS, Vanier MT, Walkley SU, Lobel P. Genetic evidence for nonredundant functional cooperativity between NPC1 and NPC2 in lipid transport. Proc Natl Acad Sci U S A. 2004;101(16):5886-91, 10.1073/pnas.0308456101.

62. Castellano BM, Thelen AM, Moldavski O, Feltes M, van der Welle REN, Mydock-McGrane L, Jiang X, van Eijkeren RJ, Davis OB, Louie SM, Perera RM, Covey DF, Nomura DK, Ory DS, Zoncu R. Lysosomal cholesterol activates mTORC1 via an SLC38A9-Niemann-Pick C1 signaling complex. Science. 2017;355(6331):1306-11, 10.1126/science.aag1417.

63. Xu J, Dang Y, Ren YR, Liu JO. Cholesterol trafficking is required for mTOR activation in endothelial cells. Proc Natl Acad Sci U S A. 2010;107(10):4764-9, 10.1073/pnas.0910872107.

64. Lu F, Liang Q, Abi-Mosleh L, Das A, De Brabander JK, Goldstein JL, Brown MS. Identification of NPC1 as the target of U18666A, an inhibitor of lysosomal cholesterol export and Ebola infection. Elife. 2015;4:e12177. 10.7554/eLife.12177.

65. Corpataux JM, Naik J, Porter KE, London NJM. The effect of six different statins on the proliferation, migration, and invasion of human smooth muscle cells. J Surg Res. 2005;129(1):52-6, 10.1016/j.jss.2005.05.016.

66. Sun W, Lee T-S, Zhu M, Gu C, Wang Y, Zhu Y, Shyy JYJ. Statins activate AMP-activated protein kinase in vitro and in vivo. Circulation. 2006;114(24):2655-62, 10.1161/CIRCULATIONAHA.106.630194.

67. Inoki K, Kim J, Guan K-L. AMPK and mTOR in cellular energy homeostasis and drug targets. Annu Rev Pharmacol Toxicol. 2012;52:381-400, 10.1146/annurev-pharmtox-010611-134537.

68. Holczer M, Hajdú B, Lőrincz T, Szarka A, Bánhegyi G, Kapuy O. A Double Negative Feedback Loop between mTORC1 and AMPK Kinases Guarantees Precise Autophagy Induction upon Cellular Stress. Int J Mol Sci. 2019;20(22):5543, 10.3390/ijms20225543.

69. Houghton PJ. Everolimus. Clin Cancer Res. 2010;16(5):1368-72, 10.1158/1078-0432.CCR-09-1314.

70. Cerni S, Shafer D, To K, Venketaraman V. Investigating the Role of Everolimus in mTOR Inhibition and Autophagy Promotion as a Potential Host-Directed Therapeutic Target in Mycobacterium tuberculosis Infection. J Clin Med. 2019;8(2):232, 10.3390/jcm8020232.

71. Jeon S-Y, Yhim H-Y, Lee N-R, Song E-K, Kwak J-Y, Yim C-Y. Everolimus-induced activation of latent Mycobacterium tuberculosis infection in a patient with metastatic renal cell carcinoma. Korean J Intern Med. 2017;32(2):365-8, 10.3904/kjim.2015.121.

72. Dara Y, Volcani D, Shah K, Shin K, Venketaraman V. Potentials of Host-Directed Therapies in Tuberculosis Management. J Clin Med. 2019;8(8):1166, 10.3390/jcm8081166.

73. Inpanathan S, Botelho RJ. The Lysosome Signaling Platform: Adapting With the Times. Front Cell Dev Biol. 2019;7:113-, 10.3389/fcell.2019.00113.

74. Napolitano G, Ballabio A. TFEB at a glance. J Cell Sci. 2016;129(13):2475-81, 10.1242/jcs.146365.

75. Brady OA, Martina JA, Puertollano R. Emerging roles for TFEB in the immune response and inflammation. Autophagy. 2018;14(2):181-9, 10.1080/15548627.2017.1313943.

76. Silwal P, Kim JK, Yuk J-M, Jo E-K. AMP-Activated Protein Kinase and Host Defense against Infection. Int J Mol Sci. 2018;19(11):3495, 10.3390/ijms19113495.

77. Jaramillo M, Gomez MA, Larsson O, Shio MT, Topisirovic I, Contreras I, Luxenburg R, Rosenfeld A, Colina R, McMaster RW, Olivier M, Costa-Mattioli M, Sonenberg N. Leishmania repression of host translation through mTOR cleavage is required for parasite survival and infection. Cell Host Microbe. 2011;9(4):331-41, 10.1016/j.chom.2011.03.008.

78. Weichhart T, Costantino G, Poglitsch M, Rosner M, Zeyda M, Stuhlmeier KM, Kolbe T, Stulnig TM, Hörl WH, Hengstschläger M, Müller M, Säemann MD. The TSC-mTOR signaling pathway regulates the innate inflammatory response. Immunity. 2008;29(4):565-77, 10.1016/j.immuni.2008.08.012.

79. Salamon H, Bruiners N, Lakehal K, Shi L, Ravi J, Yamaguchi KD, Pine R, Gennaro ML. Cutting edge: Vitamin D regulates lipid metabolism in Mycobacterium tuberculosis infection. J Immunol. 2014;193(1):30-4, 10.4049/jimmunol.1400736.

80. Yang N-C, Ho W-M, Chen Y-H, Hu M-L. A convenient one-step extraction of cellular ATP using boiling water for the luciferin-luciferase assay of ATP. Anal Biochem. 2002;306(2):323-7, 10.1006/abio.2002.5698.

81. Yamaguchi KD, Ruderman DL, Croze E, Wagner TC, Velichko S, Reder AT, Salamon H. IFN-beta-regulated genes show abnormal expression in therapy-naïve relapsing-remitting MS mononuclear cells: gene expression analysis employing all reported protein-protein interactions. J Neuroimmunol. 2008;195(1-2):116-20, 10.1016/j.jneuroim.2007.12.007.

82. Croze E, Yamaguchi KD, Knappertz V, Reder AT, Salamon H. Interferon-beta-1b-induced short- and long-term signatures of treatment activity in multiple sclerosis. Pharmacogenomics J. 2013;13(5):443-51, 10.1038/tpj.2012.27.

83. Lachmann A, Xu H, Krishnan J, Berger SI, Mazloom AR, Ma’ayan A. ChEA: transcription factor regulation inferred from integrating genome-wide ChIP-X experiments. Bioinformatics. 2010;26(19):2438-44, 10.1093/bioinformatics/btq466.

84. Zyla J, Marczyk M, Domaszewska T, Kaufmann SHE, Polanska J, Weiner J. Gene set enrichment for reproducible science: comparison of CERNO and eight other algorithms. Bioinformatics. 2019;35(24):5146-54, 10.1093/bioinformatics/btz447.

